# Brain-wide impacts of sedation on spontaneous activity and auditory processing in larval zebrafish

**DOI:** 10.1101/2024.01.29.577877

**Authors:** Itia A. Favre-Bulle, Eli Muller, Conrad Lee, Leandro A. Scholz, Joshua Arnold, Brandon Munn, Gabriel Wainstein, James M. Shine, Ethan K. Scott

**Affiliations:** Queensland Brain Institute, The University of Queensland, 4067 Brisbane, Australia; School of Mathematics and Physics, The University of Queensland, 4067 Brisbane, Australia; Brain and Mind Centre, University of Sydney, Sydney, Australia; Department of Anatomy and Physiology, University of Melbourne, 3052 Melbourne, Australia

## Abstract

Despite their widespread use, we have limited knowledge of the mechanisms by which sedatives mediate their effects on brain-wide networks. This is, in part, due to the technical challenge of observing activity across large populations of neurons in normal and sedated brains. In this study, we examined the effects of the sedative dexmedetomidine, and its antagonist atipamezole, on spontaneous brain dynamics and auditory processing in zebrafish larvae. Our brain-wide, cellular-resolution calcium imaging reveals, for the first time, the brain regions involved in these network-scale dynamics and the individual neurons that are affected within those regions. Further analysis reveals a variety of dynamic changes in the brain at baseline, including marked reductions in spontaneous activity, correlation, and variance. The reductions in activity and variance represent a “quieter” brain state during sedation, an effect that causes highly correlated evoked activity in the auditory system to stand out more than it does in un-sedated brains. We also observe a reduction in auditory response latencies across the brain during sedation, suggesting that the removal of spontaneous activity leaves the core auditory pathway free of impingement from other non-auditory information. Finally, we describe a less dynamic brain-wide network during sedation, with a higher energy barrier and a lower probability of brain state transitions during sedation. In total, our brain-wide, cellular-resolution analysis shows that sedation leads to quieter, more stable, and less dynamic brain, and that against this background, responses across the auditory processing pathway become sharper and more prominent.

**Significance Statement:** Animals’ brain states constantly fluctuate in response to their environment and context, leading to changes in perception and behavioral choices. Alterations in perception, sensorimotor gating, and behavioral selection are hallmarks of numerous neuropsychiatric disorders, but the circuit- and network-level underpinnings of these alterations are poorly understood.

Pharmacological sedation alters perception and responsiveness and provides a controlled and repeatable manipulation for studying brain states and their underlying circuitry. Here, we show that sedation of larval zebrafish with dexmedetomidine reduces brain-wide spontaneous activity and locomotion but leaves portions of brain-wide auditory processing and behavior intact. We describe and computationally model changes at the levels of individual neurons, local circuits, and brain-wide networks that lead to altered brain states and sensory processing during sedation.

## Introduction

Sedatives are extensively used in medicine both to anesthetize patients and to treat psychological conditions such as acute agitation(Levy 1996; Boyer 2009; Bosch et al. 2022). While the biochemical modes of action for these drugs are often well known(Rudolph and Antkowiak 2004; Giovannitti, Thoms, and Crawford 2015), the details of their effects on brain-wide networks, and the mechanisms by which they cause altered perception and loss of consciousness, are incompletely understood. Our understanding of these mechanisms is limited by a lack of information about activity across the brain at the level of individual neurons, and how the brain-wide networks comprising these neurons change during sedation.

As a means of addressing these open questions, we have explored the effects of the sedative dexmedetomidine (DEX), as well as its antagonist atipamezole (ATI), on spontaneous activity and auditory processing in zebrafish larvae, in which it is possible to perform brain-wide calcium imaging at cellular resolution (Ahrens et al. 2012). DEX induces sedation by decreasing activity of noradrenergic neurons in the locus coeruleus (LC)(Jorm and Stamford 1993; Guo et al. 1996). More specifically, it is a highly selective α2-adrenergic receptor (α2-AR) agonist that activates central pre- and postsynaptic inhibitory α2-receptors in the LC, thereby inducing a state of unconsciousness similar, but not identical to natural sleep (Akeju and Brown 2017; Guldenmund et al. 2017). Its effects are reversed by the α2 antagonist ATI, which blocks norepinephrine’s feedback inhibition on nociceptors, thereby competing with DEX for α2-adrenergic receptors.

In zebrafish, the administration of DEX and ATI has similar effects to those in humans(Ruuskanen, Peitsaro, et al. 2005; Maximino and Herculano 2010). The α2-AR is well conserved from zebrafish to humans(Ruuskanen, Laurila, et al. 2005), with similar ligand-binding profiles across the species(Ruuskanen, Peitsaro, et al. 2005). Behaviorally, a recent study used DEX in larval zebrafish to cause sedation and anesthesia(Bedell et al. 2020). Combined, these observations suggest that both mechanistic and behavioral effects of these drugs are conserved between zebrafish and humans.

Given zebrafish’s unique advantages among vertebrates for the study of brain-wide spontaneous activity and sensory processing networks(Vanwalleghem, Ahrens, and Scott 2018; Bedell et al. 2018), we judged that zebrafish would be a suitable model for exploring the network-level mechanisms of DEX’s and ATI’s effects on wakefulness and consciousness. Several groups have used 2-photon or light-sheet fluorescence imaging, combined with genetically-encoded calcium indicators, to image whole larval zebrafish brains, in vivo, with cellular resolution(Favre-Bulle et al. 2018; Taylor et al. 2018; Ahrens et al. 2012; Kappel et al. 2022; Tunbak et al. 2020; Cong et al. 2017; Wolf et al. 2015; Migault et al. 2018). Such studies have provided the details of individual neurons’ activity across the entire brain during spontaneous activity (Avitan et al. 2017) and during the processing of stimuli across the auditory(Constantin et al. 2020; Poulsen et al. 2021; Vanwalleghem, Heap, and Scott 2017), visual(Heap et al. 2018; Mancienne et al.; Marquez Legorreta et al. 2022; Thompson and Scott 2016; Chen et al. 2018; Dragomir, Štih, and Portugues 2020; Fotowat and Engert 2023; Karpenko et al. 2020), vestibular(Favre-Bulle et al. 2017; Favre-Bulle et al. 2020; Favre-Bulle et al. 2018; Migault et al. 2018), and water flow sensing(Vanwalleghem et al. 2020) systems.

In this study, we have revealed brain dynamics of zebrafish larvae under sedation during periods of spontaneous activity as well as in response to acoustic stimuli. As expected from the literature, our results show a reduction of spontaneous behavioral activity(Bedell et al. 2020) but also in brain-wide neural activity. These changes in baseline activity during sedation have further effects on auditory processing. While brain-wide auditory processing network appears to be essentially intact, with strong responses from neurons across regions that have previously been implicated in audition, we observed a reduction in response latencies across the auditory system during sedation. Behaviorally, escape responses occur with normal kinetics under sedation, but with an increase in the velocity and kinematic properties of the startle responses, suggesting a change in the sensorimotor gating or motor execution of this response when baseline activity is reduced during the sedated brain state.

## Materials and Methods

### Animals

High-speed behavioral recordings were performed with approval from the University of Melbourne Office of Research Ethics and Integrity (in accordance with ethics approval 2022-24987-35220-5). All other procedures were performed with approval from The University of Queensland Animal Welfare Unit (in accordance with ethics approval 2019/AE000341).

Zebrafish (*Danio rerio*) larvae, of both sexes, were maintained at 28.5 °C on a 14 hr ON/10 hr OFF light cycle. Adult fish were maintained, fed, and mated as previously described(Westerfield 2000). Experiments were carried out in larvae of the TL strain (behavioral experiments) or nacre-mutant *elavl3:H2B-GCaMP6s* larvae of the TL strain (calcium imaging experiments)(Chen et al. 2013).

### Free-swimming behavior and drug administration

Free swimming behavioral experiments were performed using custom-built behavioral rig. The rig consisted of an illumination system (infra-red LED array), an elevated arena where fish were placed in custom-made acrylic well plates (16 wells in a 4×4 grid or 7 wells in a circle, with wells that were 20 mm diameter). Acoustic stimuli were delivered using an amplifier (Dayton Audio DA30 2 × 15W Class D Bridgeable Mini Amplifier) and a speaker (Dayton Audio DAEX19CT-4 4 ohm / 5 W) affixed to the center of a circular array of wells (see extended data Figure 1.1). A high-speed camera (Ximea xib-64, model CB019MG-LX-X8G3) equipped with an EF mount adapter and a Sigma 17-70 mm f/2.8-4 DC Macro lens with an IR pass filter (750 nm) was used for imaging.

Zebrafish larvae at 6 days post-fertilization (dpf) were placed in individual wells containing 1 mL of E3 zebrafish medium. Larvae were then transferred to the behavioral rig and allowed to acclimate for 15 minutes prior to recording.

Spontaneous swimming was recorded at 50 frames per second (fps). During the recordings, 20 µL of a solution containing DEX, DMSO, and E3 was added with a pipette to bring each well to the desired concentration of 5 µM of DEX and less than 0.01% DMSO. For the control experiments, only DMSO + E3 was used.

Auditory responses were recorded at 400 fps with a 0.5 ms exposure time. Videos were recorded from −2 s to +2 s relative to stimulus onset stimulus (total of 4 s). Three sets of 5-minute recordings were made. During each recording, 10 auditory tones (1 kHz tone for 10 ms at 83 dB-SPL) were presented, with an inter-trial-interval of 30 seconds. Immediately after the first recording, 20 µL of a solution containing DEX, DMSO and E3 was added with a pipette to bring each well to a concentration of 0.05 µM of DEX and less than 0.0001% DMSO. After the introduction of 0.05 µM of DEX, we waited 5 minutes to allow for diffusion of the drug before the second recording began. Immediately after the second recording, each fish was carefully pipetted out of each well and placed in a well with fresh E3 (to eliminate contamination of DEX in third recording). A solution of 20 µL ATI, DMSO and E3 was added with a pipette to bring each well to a concentration of 5 µM of ATI and less than 0.003% DMSO. The third recording started 10 minutes later.

Concentrations of DEX and ATI were chosen based on behaviorally-effective doses described in the literature(Bedell et al. 2020).

### Extraction and analysis of free-swimming behavior with Deep-Lab-Cut (DLC)

To extract the pose of each fish automatically, we employed DeepLabCut (DLC)(Mathis et al. 2018). DLC is a deep learning model based on transfer learning, which means that the training starts using pre-trained weights from the ImageNet dataset and Residual Network model, thus avoiding the need to train a model from scratch and reducing the size of dataset required. Specifically, we adapted this toolset and applied a 50-layer ResNet network architecture and trained two pose models: a simple model with three key points (center of left and right eyes and one in the center of the swim bladder) and a more detailed one with fifteen key points:left and right eyes (2 key points per eye), swim bladder (1 key point), and tail (10 key points). Errors reported are of the Euclidean distance in pixels between the annotated positions and the positions predicted by the model.

The first model (3 key points) was trained on 82 frames from videos obtained in the same experimental conditions in a 95%/5% train/validation split. This achieved 0.88 pixel error in the training dataset and 0.77 pixel error in the validation dataset (95% train, 5% test split, reached after a total of 1,030,000 iterations).

The detailed 15-key point model was trained using 1437 frames from 23 different videos. The videos used were acquired from pilot data from other experiments in our laboratory. The videos varied in lighting conditions, resolution, and acquisition frame rate. After training using 90%/10% train/validation split, the model achieved a train error of 1.09 pixels and 1.43 pixel error in the validation dataset after 1,350,000 iterations.

The fish positions obtained from DLC were further analyzed in MATLAB using custom scripts (available on public github repository: https://github.com/ItiaFB/ZfishBeh_DLC/tree/main).

### Light-sheet calcium imaging and drug administration

Separate groups of larvae at 6dpf were used for whole-brain calcium imaging. A total of n=10 larvae were imaged for Figure 1 (n=5 for DEX and n=5 for DMSO control). Figure 3 to Figure 6 included a single group of n=7 larvae.

**Figure 1:**
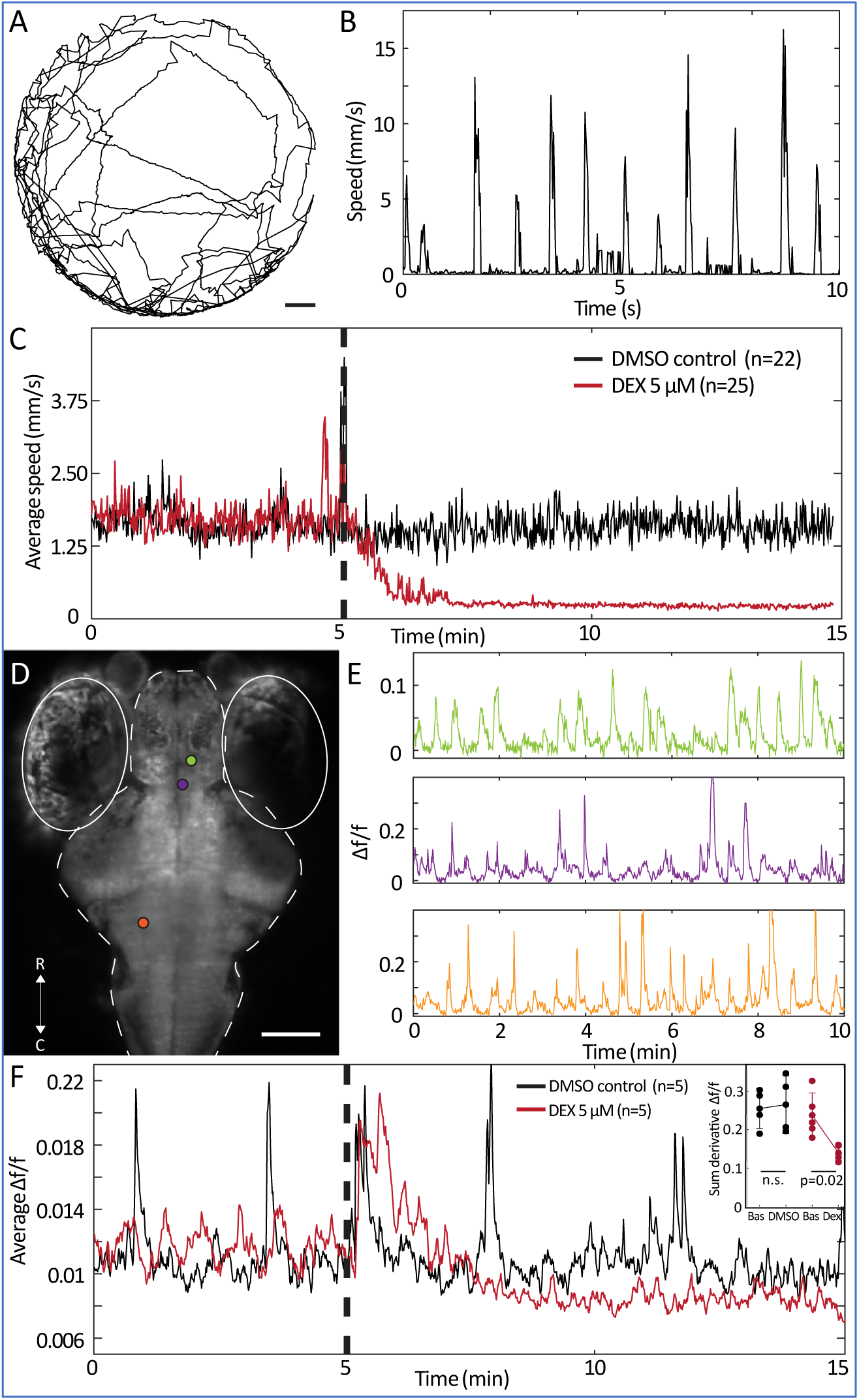
Spontaneous activity under DEX. **A.** A representative trace of a free-swimming larva during a 5-minute experiment. **B**. A representative trace of swimming speed over 10s for a free-swimming larva. **C.** Movements of larvae (averaged across 22 animals for DMSO control and 25 for DEX) for a 15-minute experiment, with the addition of DEX (red curve) or with the addition of DMSO control (black curve) at minute 5 (dashed vertical line). **D.** A maximum-projection dorsal image of a larva expressing GCaMP6s in the nuclei of all neurons. Solid lines indicate the eyes, and dashed line indicates the outline of the brain. Colored dots show the locations of ROIs with fluorescence traces shown in E. **E.** Three examples of fluorescence traces, drawn from the ROIs indicated in D. **F**. Whole brain Δf/f (averaged across 5 animals for DEX and DMSO control) for a 15-minute experiment, with the addition of DEX (red curve) or DMSO control (black curve) at minute 5 (dashed vertical line). Inset: sum of the absolute value of the first derivative of the fluorescence signal, representing fluctuations in activity between 0-5 versus 10-15 minutes in control (black, n.s. for a Friedman test) and DEX-treated animals (red, p=0.02 for a Friedman test). R: rostral; C: caudal; scale bar in A indicates 2 mm; scale bar in D indicates 100μm.

Larvae were immobilized dorsal side up in 2% low melting point agarose (Sigma-Aldrich) on microscope slides. The agarose on the rostral and left side of the fish was cut vertically, parallel to the fish, with a scalpel, and removed. The embedded fish was transferred to a 3D printed chamber(Marquez Legorreta et al. 2022). The agarose surrounding the tail was freed by removing segments of agarose perpendicular to the tail caudal to the swim bladder, and the chamber was filled with E3 media. Larvae were then transferred to a custom-built light-sheet microscope(Taylor et al. 2018) and allowed to acclimate for 15 min prior to imaging.

A mini speaker (Dayton Audio DAEX-9-4SM Skinny Mini Exciter Audio, Haptic Item Number 295-256), attached to the back wall of the chamber as previously described (Poulsen et al. 2021), was connected to an amplifier and to the computer to produce the auditory tones (84dB, 1 kHz tone for 10 ms every 30 s) for the duration of the recording.

Volumetric calcium imaging was performed as previously described(Favre-Bulle et al. 2018). An exposure time of 30 ms was chosen for each plane during volumetric imaging, with a laser power output of 60 mW, which was attenuated to 1.5 mW for each of two planes at the sample. A total Z scan of 125 μm was performed with 5 μm steps. This resulted in volumetric acquisition at 1.33 Hz. We commenced laser scanning 5 min prior to each new neural activity recording to eliminate responses to the onset of this off-target visual stimulus.

During the first set of recordings, 200 µL of DEX, DMSO and E3 was carefully added manually with a pipette to bring the chamber to 0.05 µM of DEX and less than 0.0001% DMSO. During the second set of recordings, 200 µL of ATI, DMSO and E3 was carefully added manually with a pipette to bring the chamber to 5 µM of ATI and less than 0.02% DMSO.

Recordings were done in two sets to allow the removal of DEX in between recordings. A fluidic pump (Adelab Scientific, model NE-1000X) was connected to the chamber to permit media exchanges between recordings. During each exchange, 15 mL (3 chamber volumes) of the first medium was pumped out while 15 mL of fresh E3 was delivered. A delay of approximately 10 minutes (5 min of slow water flow + 5 min of laser scanning) separated the first and second recording for each fish. Only fish that did not drift in the z-axis during the course of the whole experiment were kept for the analysis. Therefore, the same neurons within the same fish have been imaged over 30 min through baseline and exposures to DEX and ATI.

### Extraction and analysis of fluorescence traces

We used the Suite2P package to extract fluorescence traces from our raw images(Pachitariu et al. 2016). Most parameters used were default parameters. For “tau”, related to GCamp6s dynamics, we used 1.4. Other parameters related to the recordings were determined from the recording conditions. The output regions of interest (ROIs) from Suite2p, generally representing individual neurons(Vanwalleghem, Constantin, and Scott 2021), along with their corresponding fluorescent traces, were further analyzed in MATLAB.

### Registration to a reference brain

For visualization in a common reference brain, we used Advanced Normalization Tools (ANTs, https://github.com/ANTsX/ANTs) to register our results to the H2B-RFP reference of Zbrain(Avants et al. 2011; Randlett et al. 2015) as previously described(Wong et al. 2020). A set of high-resolution image volumes with 1 µm between planes was used to build a common template before registering this template to the Zbrain atlas. Motion corrected, time averaged stacks from Suite2p were then used to register individual fish to the common template and then to the Zbrain atlas as described(Marquez Legorreta et al. 2022). The resulting warps were sequentially applied to the centroids of extracted ROIs to map them all in the same frame of reference. The warped ROI coordinates were then placed in each of the 294 brain regions defined in the Zbrain atlas(Randlett et al. 2015). ROIs that were located outside of reference brain’s boundaries after registration were discarded.

### Linear regressions to acoustic stimuli

For linear regressions, regressors were built for each stimulus onset, with a typical GCaMP6s response occurring for each presentation of the stimulus [as previously described (Poulsen et al. 2021; Favre-Bulle et al. 2018)]. The coefficient of determination (r^2^) of the linear regression models was used to select stimulus responsive ROIs, and we chose a threshold based on the r^2^ distribution of our models (0.6 for high selectivity and 0.1 for low selectivity).

To gauge auditory networks at baseline and during DEX and ATI exposure, we performed linear regressions to acoustic stimuli for three 5-minute time windows: min 0 to 5 for baseline, min 10 to 15 for DEX, and min 25 to 30 for ATI.

### Selection of ROIs with reduced spontaneous activity

For the detection of ROIs with reduced spontaneous activity during DEX administration, the sum of the derivative of Δf/f of each ROI trace was calculated for baseline (0 to 5 min), DEX (10 to 15min) and ATI (25 to 30min). Values smaller than 75% from baseline and ATI to DEX were included in our analysis. A comparison of the results yielded by a range of thresholds is shown in extended data Figure 4.1.

### Energy Landscapes

To quantify dynamic modes of variance and estimate the stability of the brain states, we analyzed stimulus-locked calcium traces using an energy landscape analysis(Munn et al. 2021; Taylor et al. 2022). Briefly, we formulated an energy landscape by first computing a 1-dimensional measure of trajectories on the fluorescence timeseries data, namely the mean-squared-displacement, which is defined as

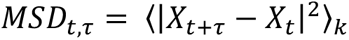

averaged over all *k* cells. The probability of observing a given MSD across the entire timeseries was then calculated using a Gaussian kernel density estimation,

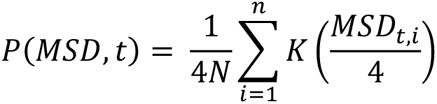

where 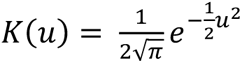. As is typical in statistical mechanics the energy of a given state, *E*_*σ*_, and its probability are related by 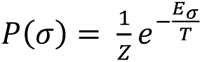 where *Z* is the normalization function and *T* is the scaling factor equivalent to temperature in thermodynamics(Tkačik et al. 2015). In our analysis ∑_*σ*_ *P*_*σ*_ = 1 → *Z* = 1 by construction and we can set *T* = 1 for the observed data. Thus, the energy (*E*) of each MSD at a given time-lag *t*, is then equal to the natural logarithm of the inverse probability, *P*(*MSD*, *t*) of its occurrence:

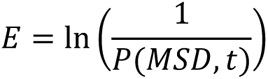

Using this approach, we analyzed dynamics in brain activity and the energies of those dynamics across space (ROIs) and time.

### Experimental Design and Statistical Analysis

Animal numbers for calcium imaging experiments were constrained by the labor intensity of the data collection methods and were n=5 for n=7 for each group, depending on the experiment (the n for each experiment is reported in figure legends and main text). Free-swimming behavioral experiments had an n of 28. As detailed elsewhere in Materials and Methods, DMSO controls were used during experiments involving the application of drugs. The statistical tests used depended on the details of the experiments and datasets, and these tests included paired t-tests, Friedman test, Wilcoxon sign-ranked tests, and Wilcoxon ranked-sum tests. The details of these statistical tests are reported in the figure legends and main text for each experiment. All statistical analyses were performed with individual animals as datapoints to avoid pseudoreplication.

## Results

### Reduction of spontaneous behavioral and brain-wide neural activity under DEX

Guided by past work(Bedell et al. 2020), we first explored the impact of DEX on spontaneous swimming behavior in zebrafish larvae. 6 dpf larvae were placed and tracked in individual wells (Figure 1A) and their spontaneous free-swimming activity was recorded over time. An example of a spontaneous trace is shown Figure 1B. Free-swimming activity was recorded before and after the administration of 5 µM DEX or DMSO control (Figure 1C). Results show a complete cessation of spontaneous activity under DEX, but no changes for DMSO controls.

We next performed brain-wide cellular-resolution calcium imaging using GCaMP6s and a custom-built light-sheet microscope, as described previously(Favre-Bulle et al. 2018). From these data, we identified regions of interest (ROIs) corresponding to individual neurons (see Materials and Methods) and used signals from these ROIs as the basis for our analyses of brain-wide activity. Examples of a fluorescence image and activity traces extracted from individual ROIs are shown in Figure 1D and E, respectively. Brain-wide fluorescence traces were recorded before and after the administration of 5 µM DEX or DMSO control (Figure 1F), and the average changes of fluorescence across all ROIs were quantified as readouts of brain-wide spontaneous activity. We found that the addition of 5 µM DEX led to, in the first instance, a burst of activity across the brain, and after 2 minutes, a decrease in average spontaneous activity below baseline and DMSO control (Figure 1C). This initial burst of activity was likely a function of water flow stimuli introduced by our addition of DEX or control solutions(Thompson et al. 2016; Vanwalleghem et al. 2020). Therefore, we restricted our further analyses to minutes 5-10 after drug administration, after water flow has ceased and DEX has taken effect. We calculated spontaneous neural activity during this interval by calculating the sum of the absolute value of the first derivative of Δf/f, which measures the fluctuations in fluorescence intensity as a proxy for activity across the brain. During this interval, spontaneous neural activity was significantly reduced in the DEX group (Fig 1F, inset), but not in DMSO control, indicating a drop in spontaneous brain activity that coincides with the cessation of spontaneous swimming behavior (Figure 1C).

### Auditory responses of larvae exposed to DEX and ATI

To explore the effects of DEX and ATI on sensorimotor gating, we recorded free swimming behavior under control, DEX, and ATI conditions while presenting auditory tap stimuli (84 dB, 1 kHz pure tone for 10 ms, every 30 s, see Materials and Methods). Responses were recorded for 5 minutes before drug treatment, followed by the application of 0.05 µM DEX for 10 minutes, and finally 5 µM of ATI for 15 minutes. Swim bouts were categorized as either forward swims or escape responses based on their location on a bimodal distribution of maximum velocities (with a threshold of 90 mm/s, Figure 2A).

**Figure 2:**
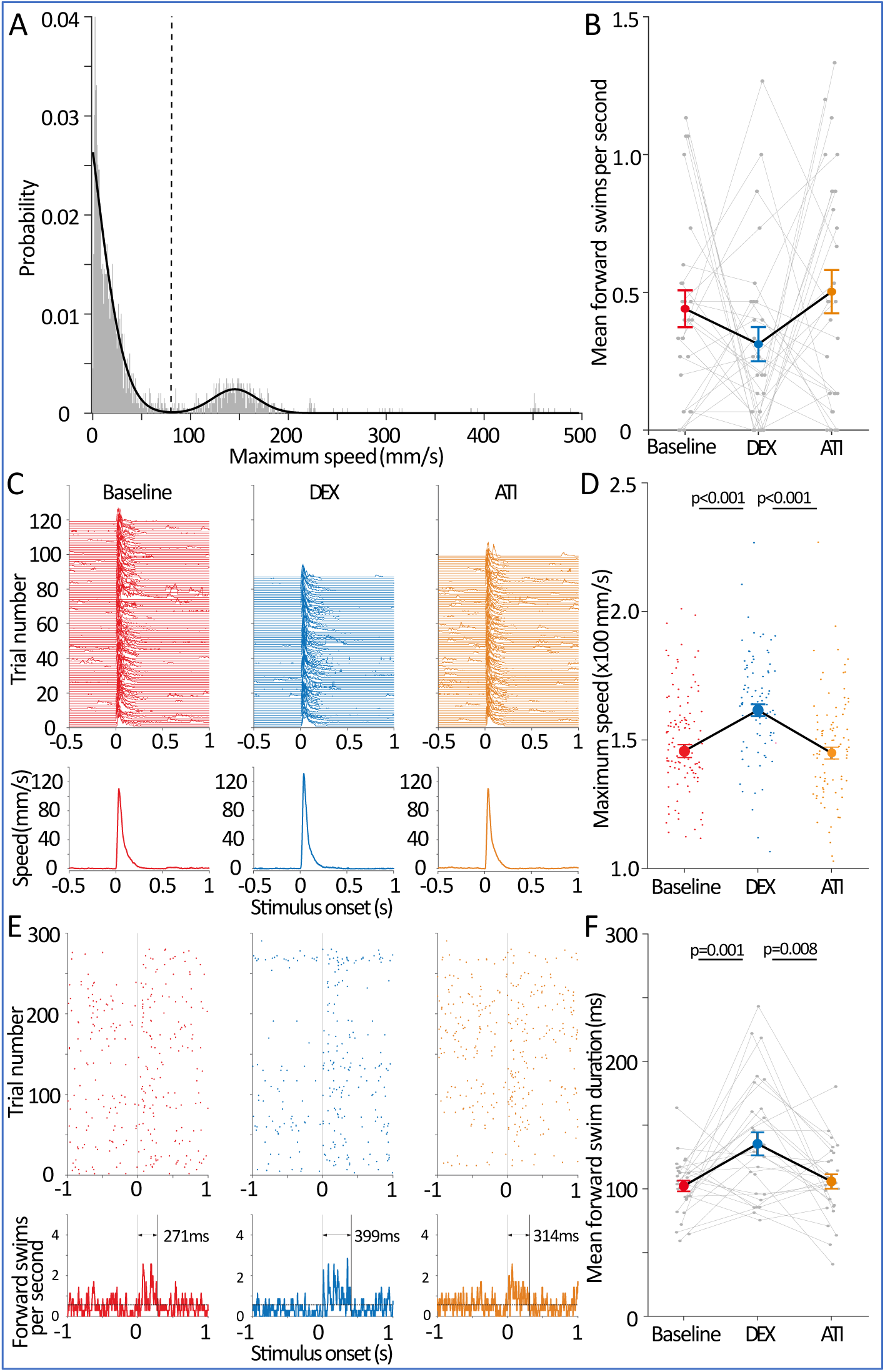
Auditory behavioral responses under DEX and ATI. **A.** Distribution of 1994 bouts (across baseline, DEX, and ATI periods), measured as each bout’s maximum speed. **B.** Average number of spontaneous forward swims, measured during the 2-second period prior to stimulus onset. **C.** Individual trials (top) and average speed (bottom) for auditory escape responses during baseline (red), DEX (blue), and ATI (orange). **D.** Distributions (individual points), means, and SEMs for maximum escape velocities in each condition. p-values are shown for Wilcoxon rank sum test. **E.** Forward swims during baseline (red), DEX (blue) and ATI (orange). Top row shows forward swim timings for all trials and bottom row shows average forward swims frequency across all trials. **F.** Forward swim duration, mean and standard error of the mean for all trials. n=28.

As expected from our results for spontaneous swimming (Figure 1), we observed a decrease in the probability of forward swim bouts during periods without acoustic stimuli (the 2s prior to stimulus onset) in DEX compared to baseline (Figure 2B, *p*<0.01, Wilcoxon signed-rank). This effect was reversed by the addition of ATI (*p*<0.01, Wilcoxon signed-rank).

In contrast, behavioral responses to acoustic stimuli remained robust. Reactions times were not significantly altered by DEX or ATI (Extended Data Figure 2.1), and in trials where escape responses occurred, the strengths of those responses, measured as peak velocity (Figure 2C), were significantly higher in DEX than at baseline or in ATI (Figure 2D, *p*<0.001, Wilcoxon ranked-sum). Next, when restricting our analysis to forward swims only (Figure 2E), we observed a significantly prolonged post-stimulus period during which forward swim responses were elevated in DEX compared to baseline (Figure 2E, *p*<0.05, Wilcoxon ranked-sum). These forward swim bouts were also significantly longer, in terms of their kinematics, than swim bouts following stimuli at baseline or in ATI (Figure 2F).

Overall, our results suggest that DEX, while it reduces or eliminates spontaneous swimming, does not interfere with behavioral auditory responses. On the contrary, responses elicited by acoustic stimuli are stronger in DEX than at baseline. In all of these regards, the application of ATI reverses the effects of DEX, returning the animals’ behavior to an approximate baseline state.

### Brain-wide auditory response dynamics in larvae under DEX and ATI

To investigate the effects of DEX and ATI on brain-wide dynamics and sensory processing, we recorded brain-wide activity while presenting acoustic stimuli (as in Figure 2). Brain-wide calcium activity was recorded in head-embedded larvae for 5 min before drug treatment as a baseline, for 10 min after the addition of 0.05 µM DEX and, finally for 15 min after the addition of 5 µM ATI. Since larvae were head-embedded, we simultaneously recorded tail movements to assure that sensorimotor transduction was intact. Consistent with our free-swimming results (Figure 2), we observed a reduction in spontaneous movements of the tail in DEX, while responses to acoustic stimuli persisted (Figure 3A; individual traces and controls are shown in Extended Data Figures 3.1 and 3.2, respectively).

**Figure 3:**
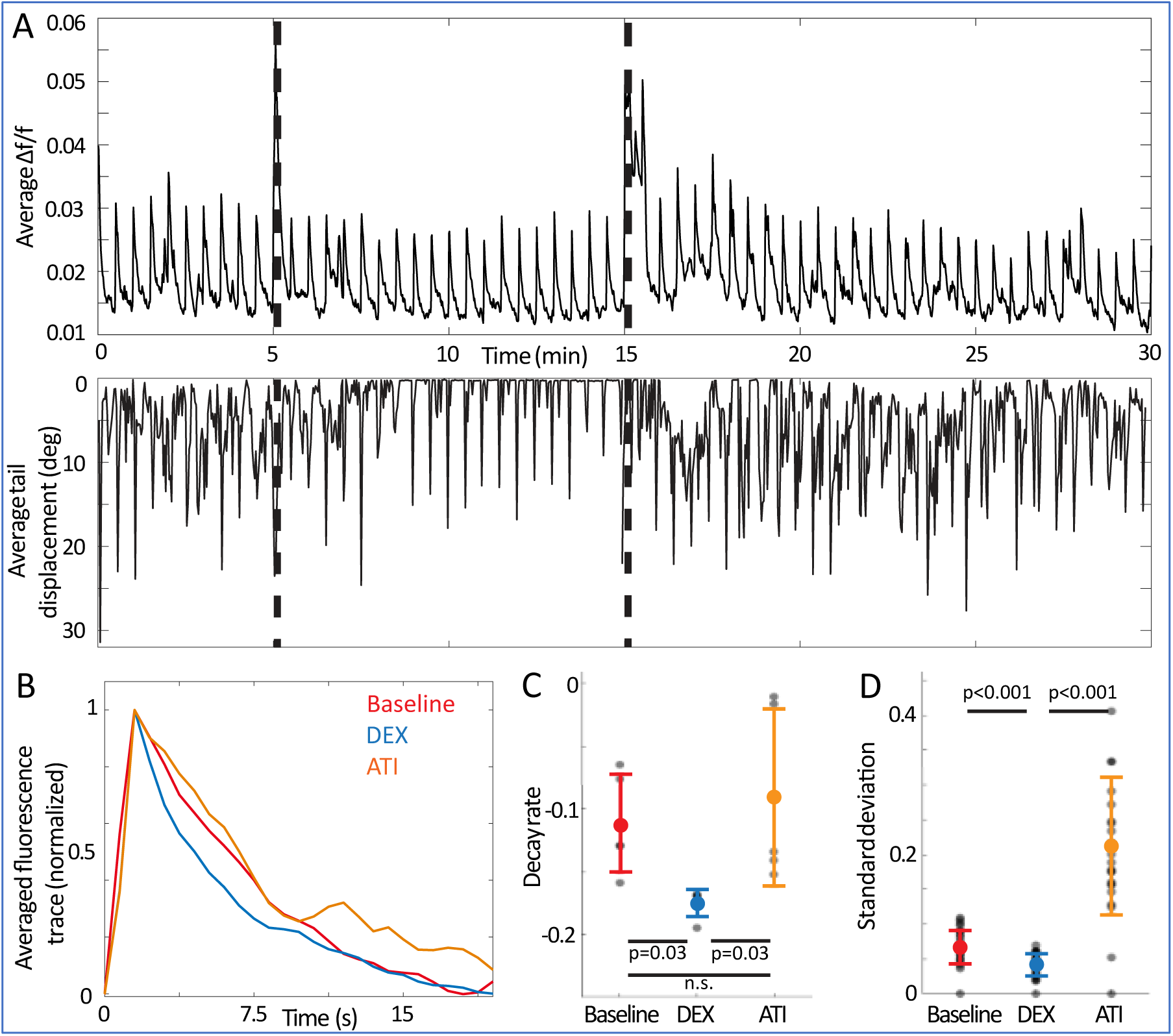
Behavioral and brain-wide responses to acoustic stimuli under DEX and ATI. **A.** Time trace of average Δf/f (top) and simultaneous average tail speed (bottom) across 7 larvae. Dashed vertical lines show the timings for DEX (5 min) and ATI (15 min) addition. Responses to acoustic stimuli are clear at 30-second intervals. **B.** Averaged fluorescence traces of whole-brain responses to acoustic stimuli during baseline (red, sampled between min 3 and 5), DEX (blue, between min 13 and 15) and ATI (orange, between min 28 and 30). **C.** Decay rates of whole-brain average responses to tones for the data shown in B (each dot indicates one animal; bars indicate mean and S.D.). p=0.009 across all three conditions (Friedman test), and with post-hoc analysis, p=0.03 (baseline vs. DEX), 0.03 (DEX vs. ATI) and 0.3 (baseline vs. ATI). **D.** Standard deviation of whole brain average response to tones for the same data. p-value is shown for a Wilcoxon signed-rank test.

Paralleling these behavioral results, calcium imaging in these animals revealed continued brain-wide responses to acoustic stimuli during DEX exposure (Figure 3A, top; individual animals’ responses and DMSO controls are shown in Extended Data Figures 3.1 and 3.2 respectively). To analyze these auditory responses in more detail, we focused on responses occurring at times of full drug effects (3-5 minutes into the experiment for baseline, 13-15 for DEX, and 28-30 for ATI). We found that neural auditory responses under the influence of DEX are sharper (with significantly faster decay rate, Figure 3B and C, Friedman test p=0.009), and more stereotyped (with significantly lower standard deviation, Figure 3D, *p*<0.001, Wilcoxon signed-rank) than during baseline, and that both of these effects are reversed with the application of ATI (Figure 2C and D).

These results suggest that DEX has multiple effects on neural activity and behavior. It reduces spontaneous neural activity and swimming, but leaves auditory activity and responses intact, and may, in fact, be leading to clearer, sharper, and less variable responses to acoustic stimuli.

### Identifying neurons with decreased activity in DEX

We have found that DEX reduces brain-wide spontaneous activity and virtually eliminates spontaneous behavior but leaves auditory processing and behavioral responses intact. To learn more about these selective alterations of neuronal activity, we set out to identify the neurons whose activity is reduced in DEX. We identified ROIs whose fluorescence traces become less dynamic in DEX (with changes in calcium activity decreased by at least 75% versus baseline and ATI, see Materials and Methods). A lower threshold of 50% yielded similar results (Extended Data Figure 4.1).

We found a large population of ROIs (almost 1,000 ROIs across 7 fish) with such reductions in activity (Figure 4A and B). These ROIs were distributed across the telencephalon, the lateral cerebellum, and in multiple rhombomeres, and this distribution was consistent across animals (Figure 4D). Based on their average responses through the experiment (Figure 4B), some of these neurons respond to acoustic stimuli during DEX treatment (the numbers and locations of these ROIs is shown in Extended Data Figure 4.2). Because of the higher spontaneous activity at baseline and in ATI, it is difficult to gauge whether they have auditory responses during these parts of the experiment. To detect possible auditory responses at baseline and in ATI, we looked at these ROIs’ activity in the Fourier domain (Figure 4C). Fourier transforms report on the presence of repetitive patterns in time, and since we are presenting acoustic stimuli every 30s (0.033 Hz), we expect to find the stimulus frequency and its harmonics (multiples of the stimulus frequency) in their Fourier domain if they are auditory responsive. The expected frequency and its harmonics are evident during DEX treatment (green arrow in Figure 4C), and although they are noisy, the primary frequency and weaker harmonics appear during baseline and ATI periods as well (examples of individual ROIs are shown in Extended Data Figure 4.3).

**Figure 4:**
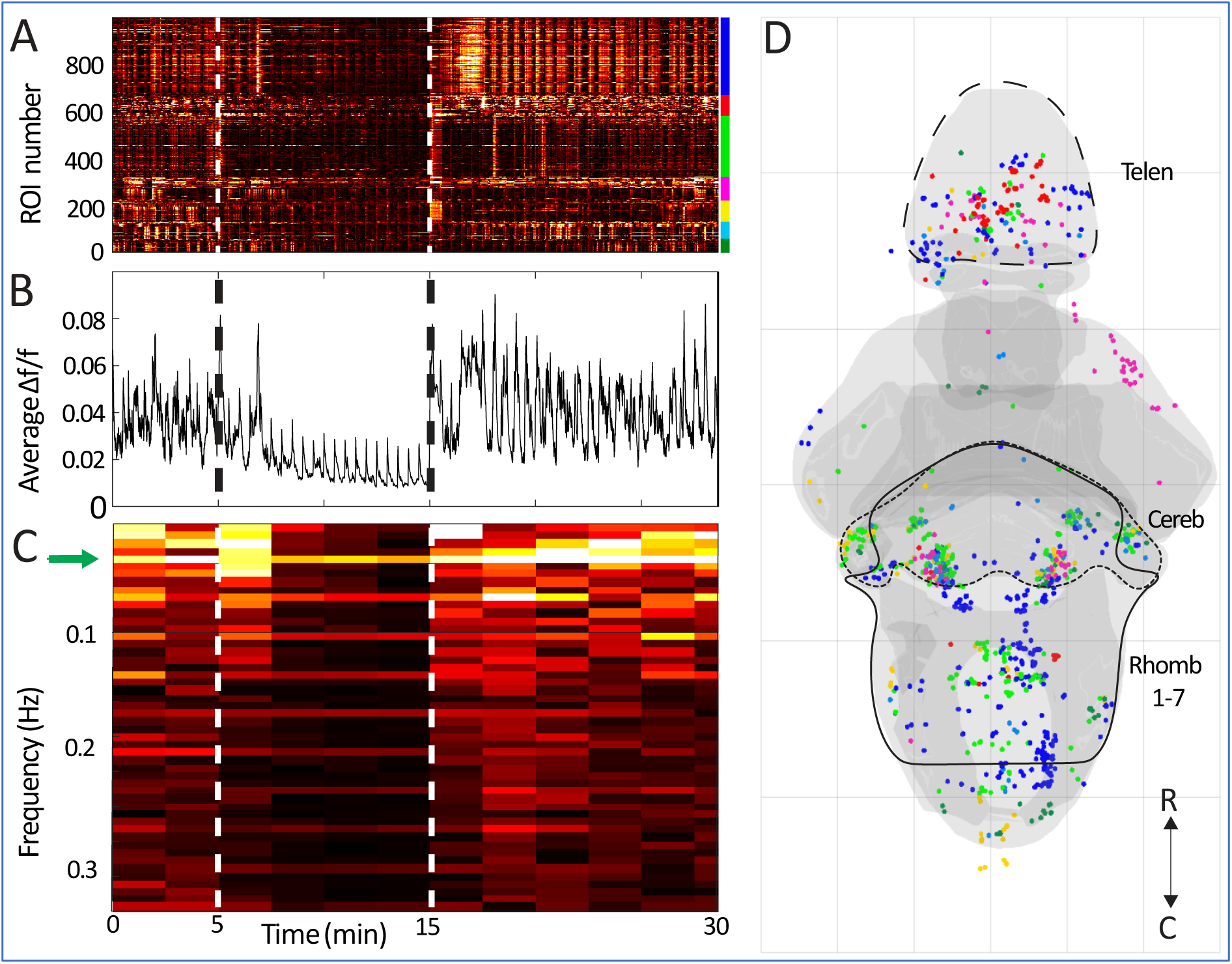
Identification of ROIs with reduced activity under sedation. **A.** Raster plot of all ROIs with at least 75% decrease in fluorescence activity when exposed to DEX (n=7 larvae). Color bar (right) indicates the ROIs belonging to each of seven larvae. **B.** Average Δf/f through time of all of the ROIs fluorescence traces shown in A. **C.** Fourier transform of average Δf/f time trace (in B) for different time windows (2.5 min in duration) across the timeline. **D.** Spatial distribution of ROIs with reduced activity during DEX exposure (from A). Colors indicate the larva that each ROI was drawn from (colors from A, right). Dashed vertical lines show the timings for DEX (5 min) and ATI (15 min) addition. R: rostral; C: caudal.

These results reinforce the idea that DEX’s primary impact on the auditory network is a reduction in activity at times when acoustic stimuli are not present. The effect, especially in neurons whose spontaneous activity is most dramatically reduced, is to elevate the prominence of the auditory responses that these neurons have. This, in turn, has implications for how the brain-wide processing of acoustic stimuli may occur in DEX, and may explain some of the effects of DEX that we have seen on the animals’ auditory behavior.

### Auditory networks at baseline and during sedation

To identify the neurons composing the auditory processing network, we performed a linear regression between the acoustic stimuli and each ROI’s activity (Bzdok and Ioannidis 2019). In this analysis, ROIs whose responses are strongly correlated with stimuli have high r^2^ values and are deemed to be involved in the perception or processing of the stimulus. We found that the distribution of r^2^ values was shifted toward higher values during DEX treatment (Figure 5A), indicating stronger correlations between the stimuli and the responses of ROIs across the brain as compared to baseline and ATI conditions.

**Figure 5:**
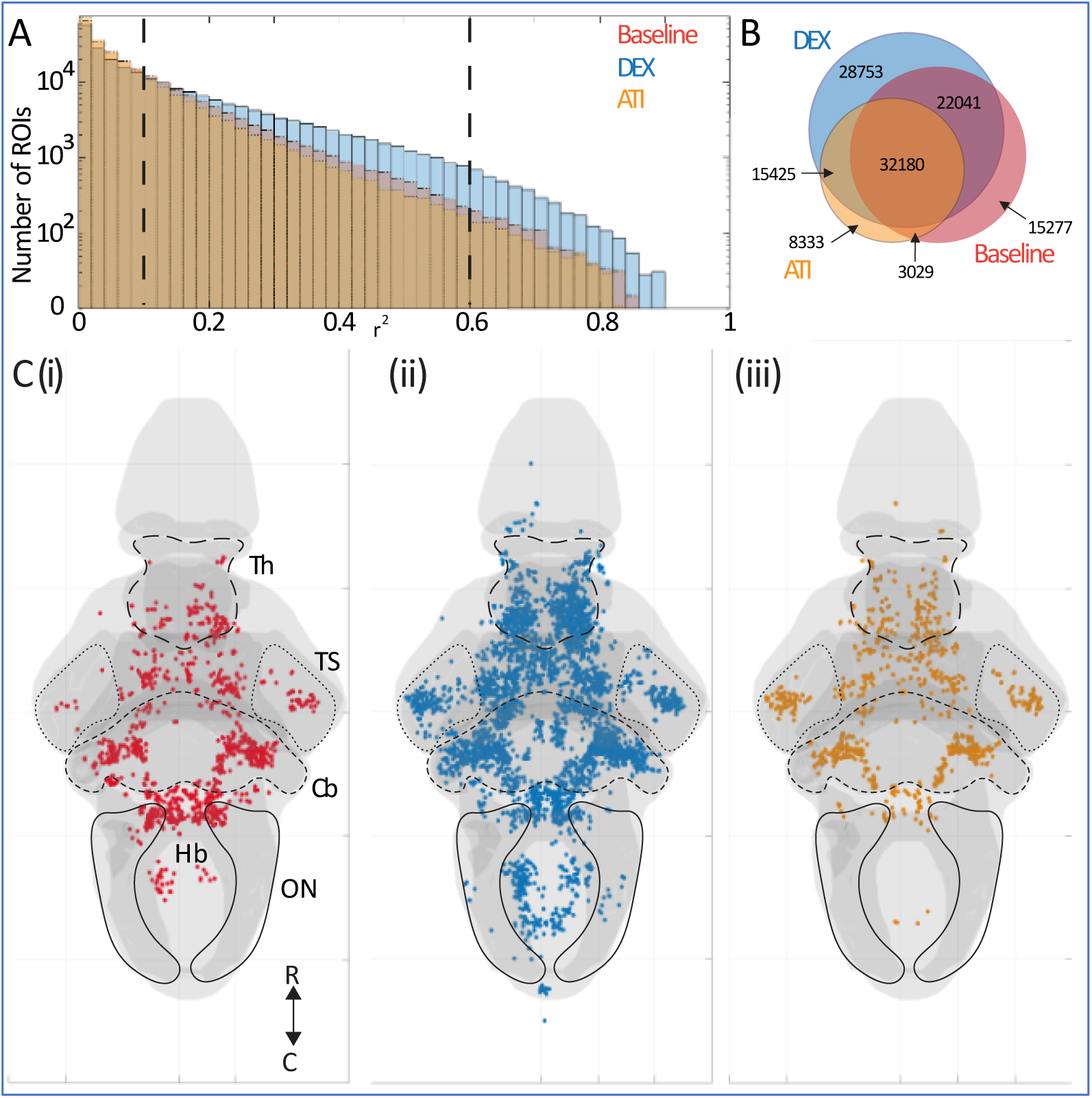
Detection of brain-wide auditory networks. **A.** Distributions of r^2^ values for all ROIs’ responses to acoustic stimuli (red, baseline; blue, DEX; orange, ATI). Dashed vertical lines show threshold of 0.1 and 0.6 used for panels B and C respectively. **B.** Venn diagram showing the overlap of auditory ROIs from A at baseline, in DEX, and in ATI at low selectivity r^2^>0.1. **C.** Spatial distribution of auditory ROIs from A at (**i**) baseline, (**ii**) in DEX and (**iii**) in ATI at high selectivity r^2^>0.6 for clarity. Auditory responses are prominent in the octavolateralis nucleus (ON), torus semicircularis (TS), Thalamus (Th), Cerebellum (Cb), and the remainder of the hindbrain (Hb). n=7 larvae. R: rostral; C: caudal.

Using both low (0.1) and high (0.6) r^2^ thresholds to define auditory responsive ROIs (see Extended Data Figure 5.1 for results using intermediate thresholds), we compared the numbers and distributions of neurons identified in the auditory system in each of the three conditions (Figure 5B for r^2^>0.1). Auditory ROIs in DEX include most of the ROIs identified as auditory during baseline (75% for low selectivity r^2^>0.1, 51% for r^2^>0.6) and ATI (81% for low selectivity r^2^>0.1, 77% for r^2^>0.6). However, the reciprocal is not true; more than 45% of ROIs identified as auditory during DEX are not identified as auditory at baseline or in ATI using low selectivity r^2^>0.1, a figure that rises to 80% when high selectivity of r^2^>0.6 is applied. While spatial distributions of these ROIs (Figure 5C) show a greater number of ROIs identified as auditory during DEX, this appears to be an essentially uniform increase across auditory brain regions [notably the octavolateralis nucleus (ON), the torus semicircularis (TS), the thalamus (Th), the cerebellum (Cb), and the remainder of the hindbrain (Hb)]. Both the overlap in auditory ROIs (Figure 5B) and the locations of these ROIs (Figure 5C) are consistent with the idea that, in DEX, the core auditory system is expanded to include additional neurons, but these results do not point to any particular functionally or anatomically distinct population of neurons that join the auditory network as a result of sedation.

In total, the results from our behavioral and brain imaging experiments suggest that DEX reduces baseline activity in the brain without interfering with the processing of or response to acoustic stimuli. The reduction in spontaneous neural activity caused by DEX leads to 1) reduced spontaneous swimming, and 2) increased salience of auditory responses in the brain, since they are occurring over a quieter baseline. This increased salience appears to be the reason for our detecting more auditory responses in DEX than at baseline, and this effect appears to be roughly evenly distributed across the auditory network. All of these effects are reversed when ATI is applied, suggesting an adrenergic mechanism for the behavioral and network effects that we observe.

### Temporal properties of auditory responses in DEX and ATI

In terms of the network-scale mechanisms, it is not immediately evident how DEX strongly suppresses spontaneous activity but permits sensory processing. It is also not clear why the strengths of behavioral responses are increased in DEX. In order to better understand whole-brain dynamics, we next explored the temporal properties of auditory processing at baseline, in DEX, and in ATI, and described the relative stability of brain-wide networks under each of these conditions.

We used a cross-correlation function to explore auditory response dynamics in greater temporal detail. Specifically, we compared the signals through time for each stimulus event to an ideal GCaMP6s response function and found the temporal lag that maximizes this correlation for each ROI (Figure 6A). Each ROI’s response delay (or lag, given in number of successive measurements or frames) was thereby calculated for each stimulus event. We generated distributions of the response lags for ROIs across the brain at baseline and in DEX and ATI (Figure 6B). We found more stereotyped auditory responses at smaller lags (Δt=0 to 1 frame, equal to 0 to 0.75 s) in the DEX condition than at baseline or in ATI, which have more heterogeneous and less synchronous responses shifted toward longer lags (peak at Δt=2 frames, equal to 1.5 s).

**Figure 6:**
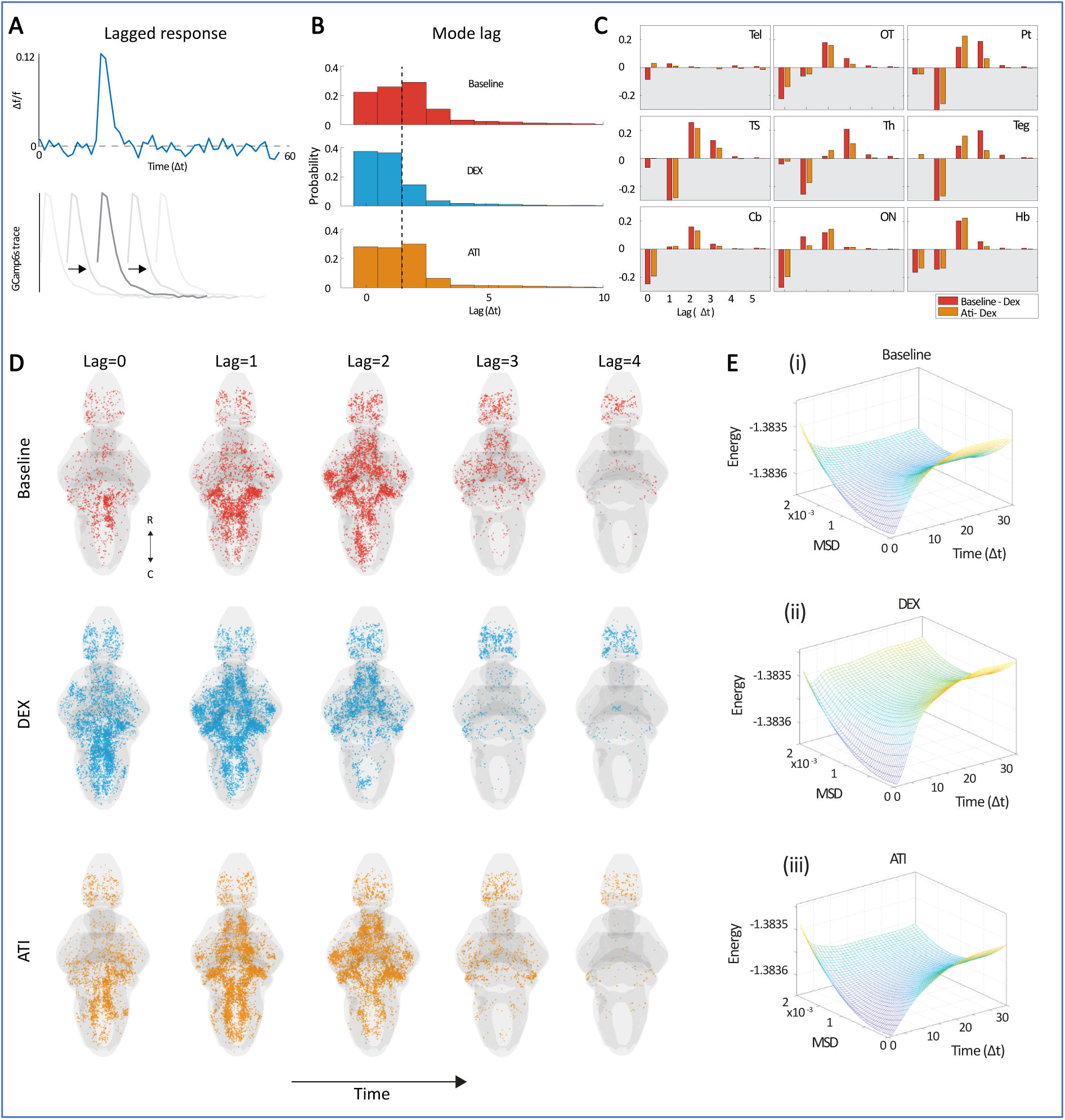
Lagged calcium responses. **A**. Example of lagged response (top). Example of GCamp6s trace moved to different onset times for cross correlation with lagged response (bottom), with the darker color indicating a stronger cross correlation. **B**. Histogram of the lags at which the peak cross-correlation values occurred, calculated across all stimulus events in whole brains, for baseline (red), DEX (blue), and ATI (orange) across 7 fish. **C**. Region-by-region differences in lags at which the cross-correlation peaked for DEX versus baseline and ATI. Negative values indicate higher relative proportions of ROIs at a given lag for the DEX treatment versus baseline (red) or ATI (orange). Tel: telencephalon; OT: Optic tectum; Pt: pretectum; TS: torus semicircularis; Th: Thalamus; Teg: tegmentum; Cb: cerebellum; ON: octavolateralis nucleus; Hb: hindbrain. **D**. Spatiotemporal maps of lagged evoked calcium response for baseline, DEX, and ATI (r > 0.95 for visualization). **E.** Energy (logarithm of inverse state probability of mean-squared displacement distribution, see Materials and Methods) against mean-squared displacement (MSD) of calcium traces and consecutive frame (Δt), across all fish. This represents the dynamic stability of brain activity across the 3 conditions: baseline **(i)** DEX **(ii)** and ATI **(iii)**. R: rostral; C: caudal.

By restricting analyses to the ROIs belonging to specific auditory brain regions, we gauged how these lags are manifested in different parts of the auditory network (Figure 6C). While responses in the telencephalon seem consistent across different lag values, in all other brain regions studied, responses in DEX occurred earlier than the equivalent responses at baseline or in ATI. This is represented as negative values for (baseline-DEX) and (ATI-DEX) at short lags and positive values at longer lags (Figure 6C). While the temporal details of these quicker responses are slightly different across regions, there is a consistent preponderance of lower-lag responses in DEX, and longer lag responses both at baseline and in ATI. These results suggest that this brain-wide auditory network processes information more quickly when DEX is present.

Viewing the spatial distribution of all ROIs for all three conditions at each lag (Figure 6D), we can see the spatiotemporal distribution of auditory responses, illustrating both the earlier occurrence of responses in DEX (consistent with Figure 6B: 1 to 2 frames, representing 0.75 to 1.5 s) and the brain regions in which these temporal effects are most dramatic (consistent with Fig 6C: all auditory brain regions except for the telencephalon). Further reflecting faster kinetics, the brain-wide correlations return to a near-baseline state after 3 frames (2.25 s) for DEX, while this takes roughly 4 frames (3 s) for baseline and ATI. We note that the telencephalon has very high spontaneous activity under normal conditions (Vargas, Þorsteinsson, and Karlsson 2012), leading to high r^2^ values that may not be true auditory responses for a random subset of telencephalic ROIs. This could explain both why this region remains active after other regions have fallen quiet (Figure 6D) and why this region’s lags do not shift across our treatments (Figure 6C).

Finally, we looked at the moment-to-moment stability of whole brain dynamics by measuring brain state energy transitions (Figure 6E, see Materials and Methods). To do so, we calculated the likelihood that neuronal activity (i.e., the brain state) would change by a prespecified amount in a specific window of time (Munn et al. 2021), see Materials and Methods). If arousal is thought to increase the general excitability of the brain, it is expected that the brain will change less through time during sedation as compared to baseline and ATI conditions. Low-probability brain state transitions are associated with a high energy barrier, whereas higher-probability state transitions are associated with a lower energy barrier. Using all timeseries from all 7 fish, we found that the energy landscape under DEX was initially deeper relative to baseline, suggesting a stable attractor state (Figure 6Ei), whereas baseline and ATI were shallow, reflecting a more unstable and temporally variable set of state transitions (Figure 6Eii-iii). DEX exposure leads to a deeper energy landscape, a more stable attractor state, and a lower probability of state transitions.

## Discussion

We have found that the application of the sedative DEX, which indirectly inhibits noradrenergic neurons in the LC, leads to a brain-wide suppression of spontaneous activity, a deepened network-wide energy landscape that makes state transitions less likely, and a cessation of spontaneous swimming behavior. All of these effects are reversed by the application of ATI, which leads to the restoration of noradrenergic signaling. We observe that the loss of spontaneous brain activity, which we presume to be the root cause of the other network and behavioral effects, does not prevent elicited activity in the brain. Indeed, responses to acoustic stimuli become sharper and more widespread in the brain-wide auditory network, and stronger behaviorally, when DEX is applied.

These relationships among spontaneous brain activity, sensory processing, and behavior, have important implications for how animals make context-dependent decisions. An alert animal navigating its environment must take its surroundings and recent experience into account when making behavioral decisions. Numerous external and internal factors impinge on sensory processing and sensorimotor gating. These range from simple mechanisms for sensorimotor gating, such as pre-pulse inhibition (PPI) and habituation, to more general context, including the presence of predators, social factors, and general arousal (Pereira and Moita 2016). We propose that many of these contextual features are encoded, at least at short time intervals, in the network’s spontaneous activity. Our results using DEX suggest that, with the loss of baseline activity in the brain, the auditory processing pathway functions unimpeded by contextual considerations, resulting in stronger, broader, and sharper activity in response to acoustic stimuli, and in more dramatic behavioral responses. It would be interesting in future work to explore which types of sensorimotor gating are impacted by DEX. For instance, it is possible that PPI is fundamental to the auditory processing pathway, and would therefore still take place under DEX, while the effects of general arousal would depend on baseline activity, and therefore be lost during sedation.

The current results also show how the brain-wide auditory network changes with the removal of its noradrenergic components, especially from the LC. Specifically, we find that neurons whose spontaneous activity is most reduced by DEX are concentrated in the lateral cerebellum and elsewhere in the hindbrain, including in regions downstream of the LC (Wang, Wang, and Mu 2022). Unsurprisingly, these regions containing neurons with reduced spontaneous activity (Figure 4) also show an increase in neurons strongly correlated to acoustic stimuli (Figure 5) in DEX. Caudal regions of the hindbrain, where premotor neurons are abundant, show a particular enrichment of auditory responsiveness in DEX, consistent with stronger sensory transmission and reduced sensorimotor gating when noradrenergic signaling from the LC is inhibited.

Notably, the reversal of this blockade with ATI restores normal behavior and brain activity according to all of the analyses that we have applied. These reversals include a return to normal spontaneous swimming (Figure 2) and spontaneous activity in the brain (Figures 2 and 4), and also a return to baseline values for acoustic escape responses and the decay rates of brain-wide auditory processing (Figure 3). Closer analyses of the auditory network reveal that similar numbers of neurons with similar correlations to acoustic stimuli are observed at baseline and after the reversal of blockade with ATI, and importantly, that ATI restores a brain-wide distribution of auditory neurons that is spatially indistinguishable from the baseline distribution (Figure 5). All of these results suggest that ATI, by releasing the blockade on noradrenergic signaling from the LC, precisely reverses the effects of DEX, rather than introducing a separate compensatory effect to restore behavior.

Looking more closely at the temporal properties of auditory responses, we observe that auditory processing is initiated at shorter lags, and that it is abbreviated, under DEX exposure (Figure 6). These effects occur on the timescale of seconds, which is too long [even accounting for GCaMP kinetics (Chen et al. 2013)] to be directly involved in the generation of a behavioral responses that typically occur in tens of milliseconds (Burgess and Granato 2007; Jain et al. 2018; Zeddies and Fay 2005; McClenahan, Troup, and Scott 2012). We speculate that extended activity in the auditory network may play a role in establishing context for future decision making. Future tests of this hypothesis could involve rapid repetitive acoustic stimulation in the presence of DEX to see whether habituation occurs during sedation. It would also be interesting to see whether recall of a habituated state (established during sedation) is possible after sedation is lifted. These tests would establish whether the neural mechanisms for habituation rest in the auditory processing pathway itself, or whether they also depend on the formation of novel patterns of activity elsewhere in the brain. Overall, the data presented here provide a tantalizing glimpse of the connections that exist among baseline brain activity, sensory processing, ethological context, and behavior, leaving many mechanistic questions open to future exploration.

Our identification of auditory ROIs depended on their correlations to the acoustic stimuli (see Material and Methods), and on this basis, we found a greater number of auditory ROIs during sedation (Figure 5). The simplest explanation for this effect is that the removal of background activity led to higher correlations for all ROIs with roles in auditory processing. Based on their distributions and response profiles, there is no widespread evidence that new ROIs are recruited to the auditory network during sedation, but rather that their responses became more salient in the absence of spontaneous activity, allowing new neurons to exceed the r^2^ threshold for inclusion as auditory. This interpretation is bolstered by our observations of activity in the Fourier domain (Figure 4), in which temporal patterns of activity that are clear during sedation are nonetheless present (in a noisier form) at baseline and in ATI. As such, while sedation is an unnatural state, it is one in which sensory processing can be detected more sensitively, and where subtle network activity that would normally be missed might become discoverable. As an example from the current study, we observed auditory ROIs in the Raphe nucleus under DEX exposure, but not at baseline or in ATI (Figure 5C). The Raphe has not been identified as part of the larval zebrafish auditory system in previous whole-brain analyses (Poulsen et al. 2021; Vanwalleghem, Heap, and Scott 2017), but past literature focusing on the Raphe has shown that it modulates sensory information(Yokogawa, Hannan, and Burgess 2012) and is activated during acoustic stimuli (Pantoja et al. 2016), potentially giving it an important role in auditory processing or decision making. While the contributions of such circuit elements would eventually have to be described in a natural, unsedated state, their initial identification could be aided by observing sensory processing under sedation. This use of a sedated state for the initial description of sensory networks, especially when using regression-based approaches and optical physiology, could be applied across sensory modalities and model systems.

## Acknowledgments

We thank the Scott and Shine labs for feedback on the manuscript. Support was provided by the Australian Research Council (DECRA DE230100972 to I.A.F, Discovery Project DP220103812 to I.A.F and E.K.S., Discovery Projects DP200102885 and DP230102614 to E.K.S and DECRA DE229199691 to C.C.Y.L), the Australian National Health and Medical Research Council (Ideas Grant 2012140 to I.A.F.), a Simons Foundation Research Award (625793) to E.K.S, the Viertel/Bellberry fellowship to J.M.S and the NHMRC (1193857) to J.M.S. The research reported in this publication was supported by the National Institute of Neurological Disorders and Stroke of the National Institutes of Health under Award Number R01NS118406 to E.K.S. The content is solely the responsibility of the authors and does not necessarily represent the official views of the National Institutes of Health. Support was also provided by the QBI Fabrication Facility.

## Extended Data Figures

**Figure 1.1:**
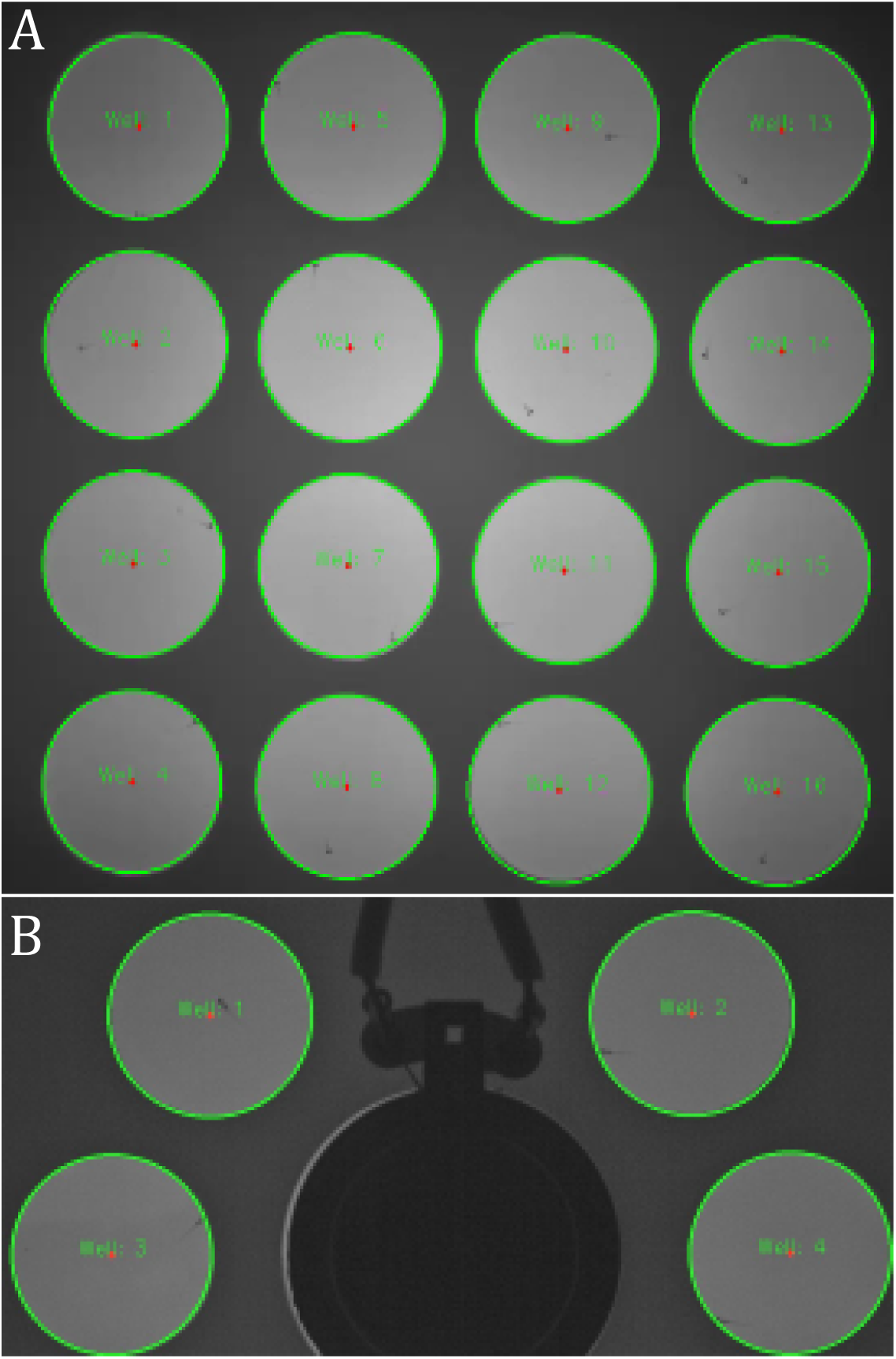
Well configuration for free-swimming experiments. **A.** 4×4 individual wells (20 mm diameter) with free swimming larvae. **B.** 4 individual wells (20 mm diameter) placed in a half circle with free swimming larvae. The speaker can be seen in the center between the wells. The wells’ centers (red dots) and borders (green outline) are automatically detected.

**Figure 2.1:**
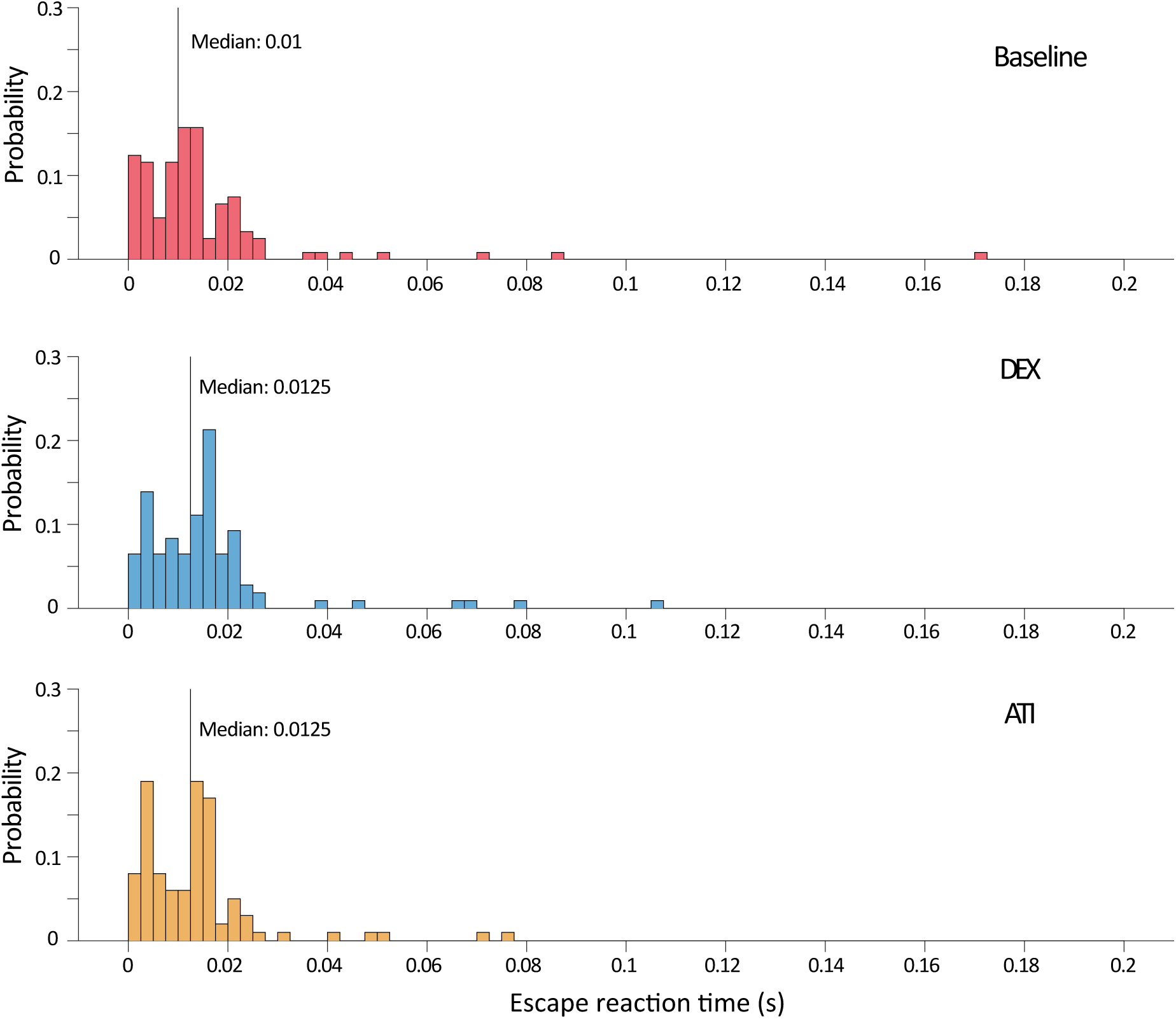
Reaction time distribution for baseline (top), DEX (centre) and ATI (bottom). Distributions and median values are indicated.

**Figure 3.1:**
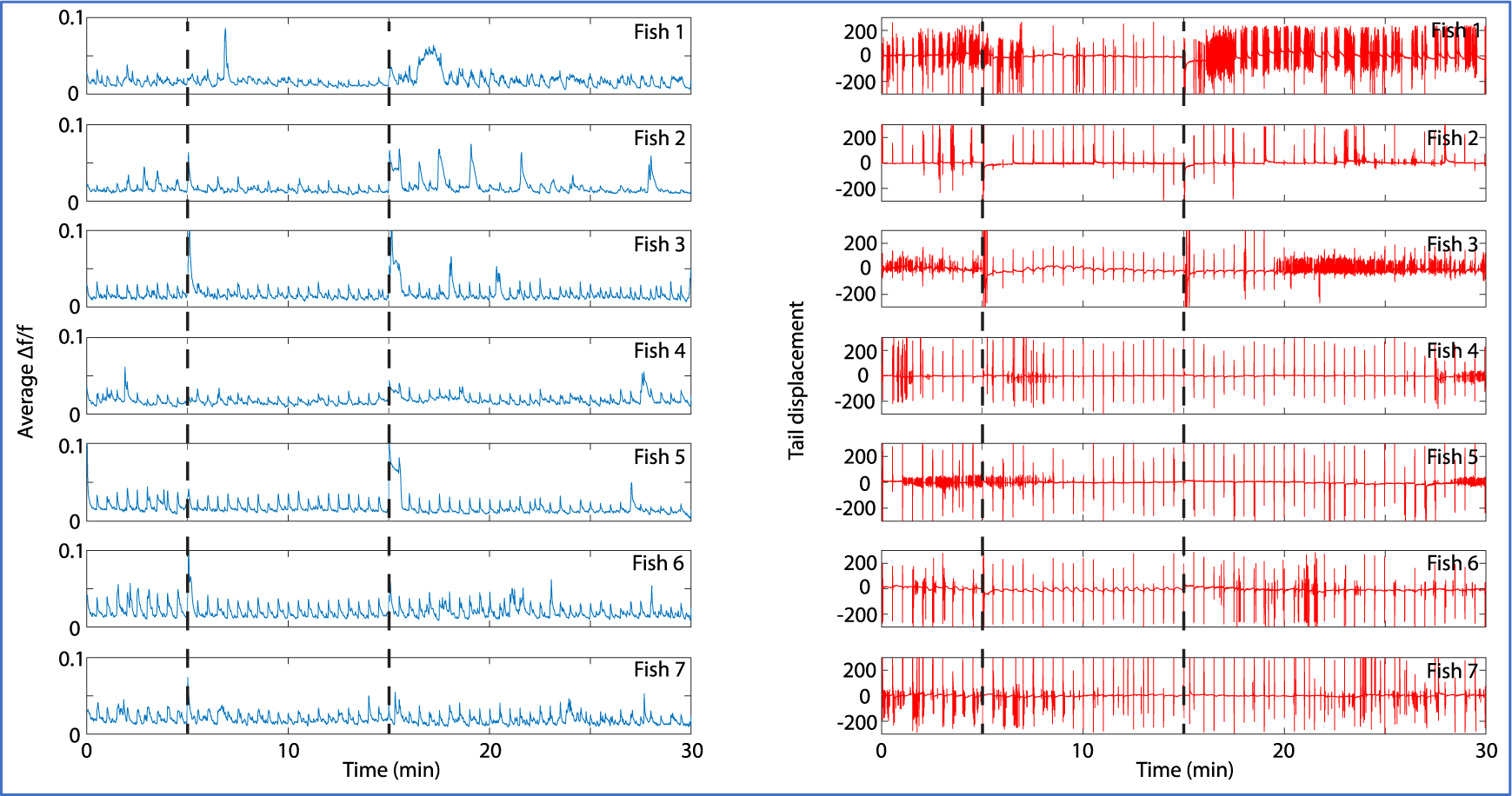
Individual brain activity and tail responses under DEX and ATI exposure. Left column shows average Δf/f over time of all ROIs in each fish undergoing DEX and ATI exposure. Right column shows tail movement for each fish over time. Vertical dashed lines show times where DEX (min 5) and ATI (min 15) were added.

**Figure 3.2:**
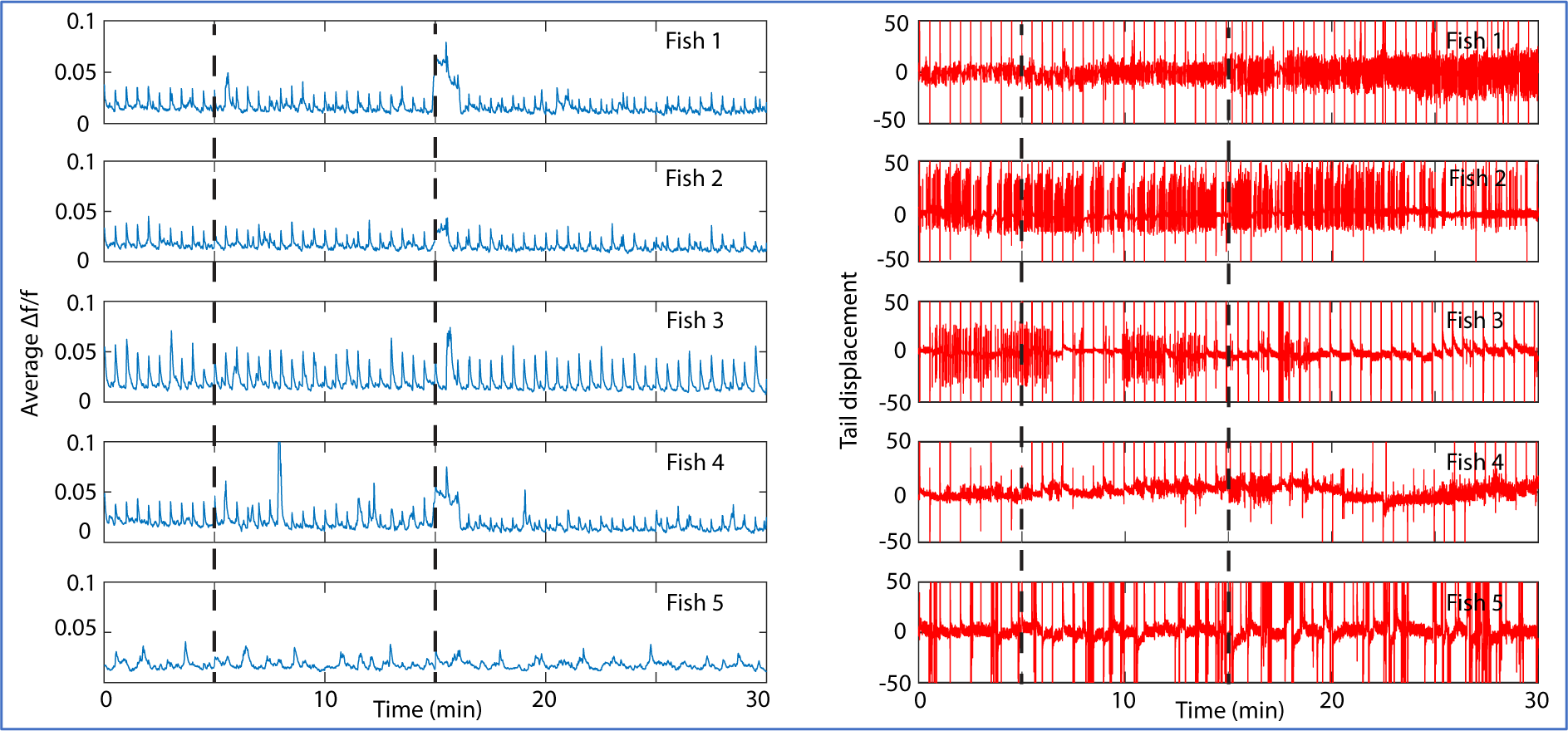
Individual brain activity and tail responses with DMSO control. Left column shows average Δf/f over time of all ROIs in each fish undergoing DMSO control. Right column shows the respective tail movement for each fish over time. Bottom line is the average across fish.

**Figure 4.1:**
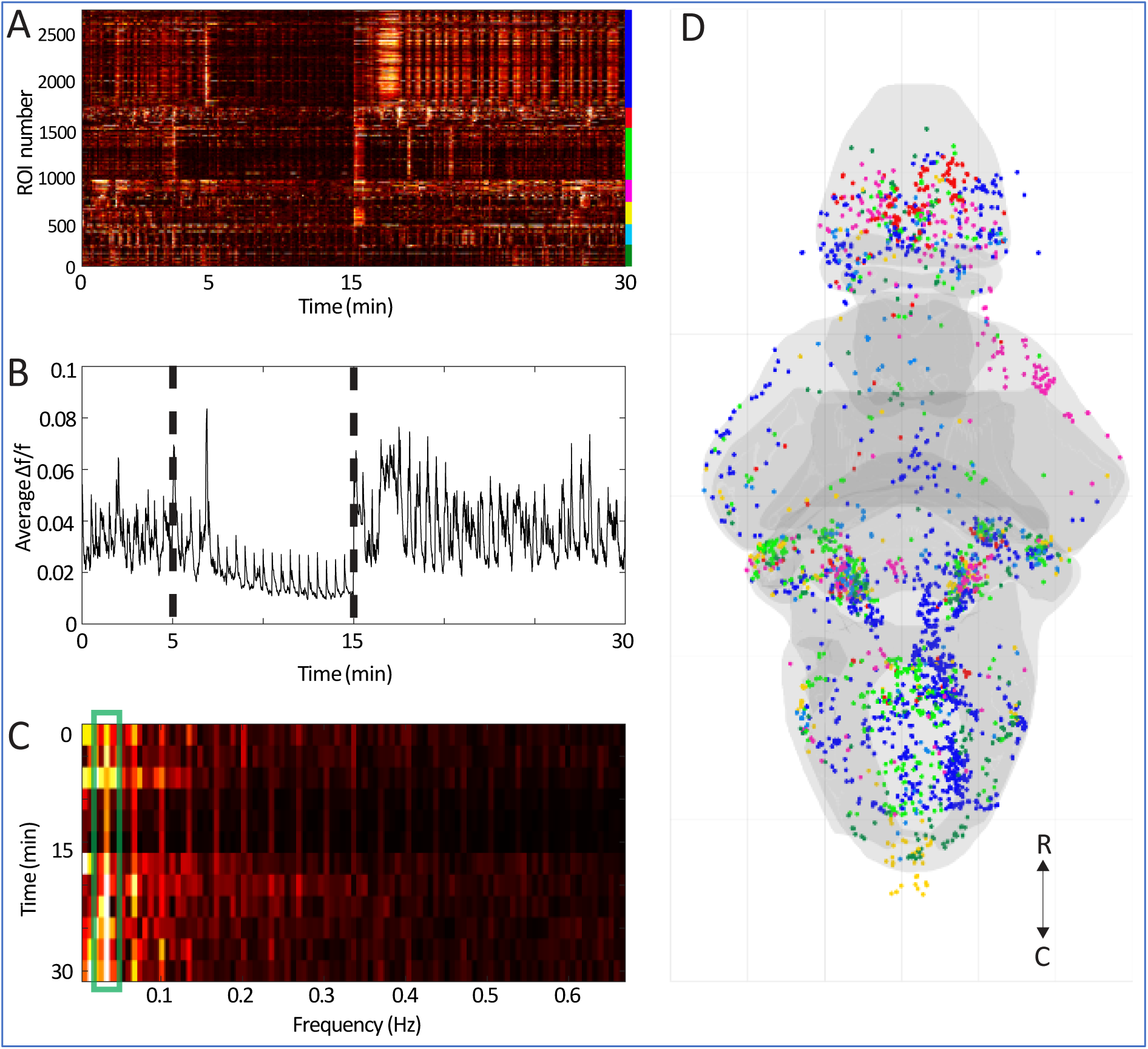
Identification of ROIs with reduced activity under sedation. **A.** Raster plot of all ROIs with at least 50% decrease in activity when exposed to DEX (n=7 larvae). Color bar (right) indicates the ROIs belonging to each of seven larvae. **B.** Average Δf/f through time for all ROIs represnted in A. **C**. Fourier transform of average Δf/f time trace (in B) for different time windows (2.5 min in duration) across the timeline. **D**. Spatial distribution of ROIs with reduced activity during DEX exposure (from A). Colors indicate the larva that each ROI was drawn from (colors from A, right). Dashed vertical lines show the timings for DEX (5 min) and ATI (15 min) addition. R: rostral; C: caudal.

**Figure 4.2:**
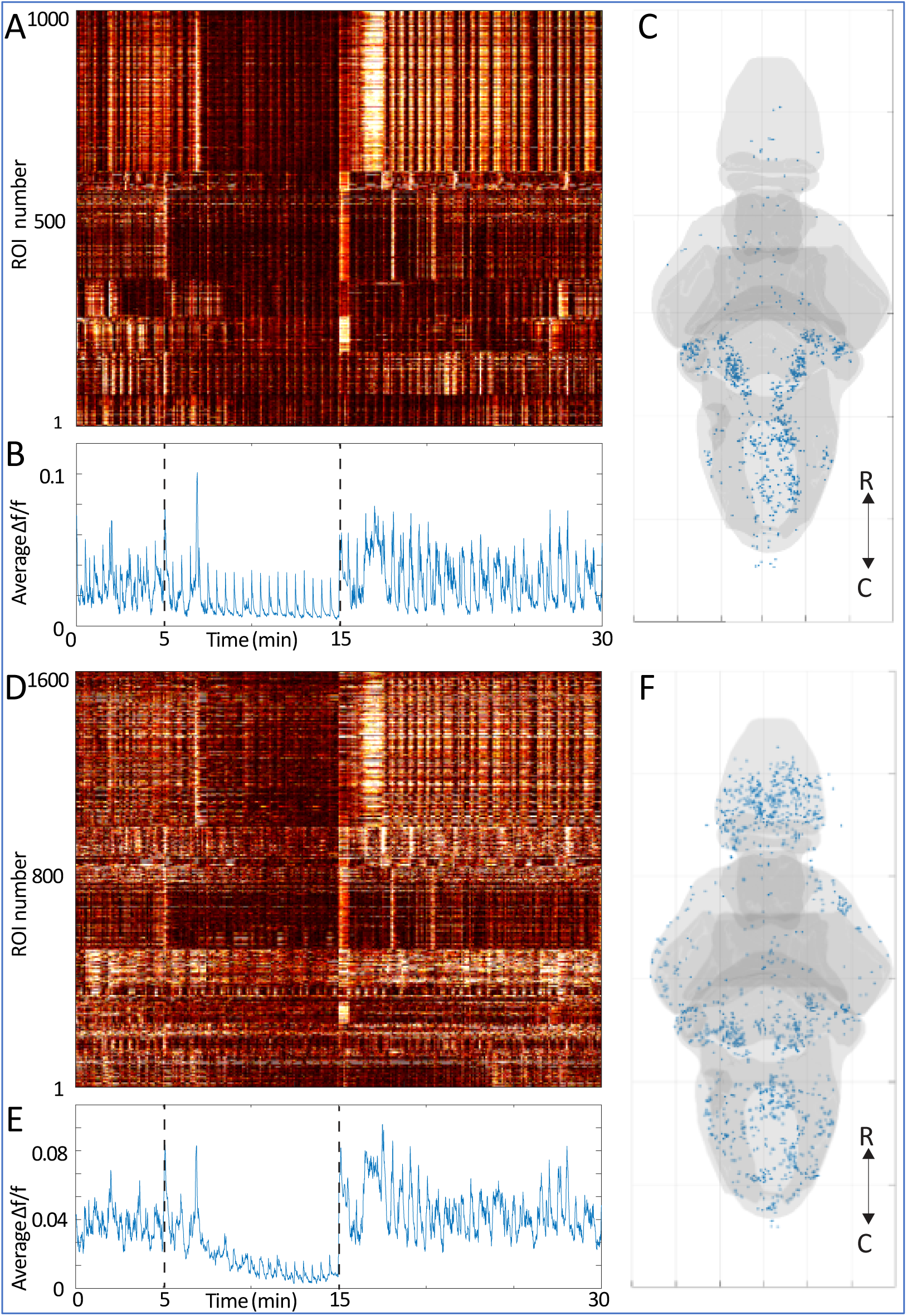
Auditory and non-auditory ROIs with responses suppressed by DEX. **A.** A Δf/f heatmap for all ROIs characterized as auditory (r2>0.1 after linear regression), among the ROIs shown in Figure 4. **B.** Average Δf/f trace for the ROIs in A. **C.** Locations of the ROIs represented in A. **D.** A Δf/f heatmap for all ROIs characterized as non-auditory (r2<0.1 after linear regression), among the ROIs shown in Figure 4. **E.** Average Δf/f for the ROIs in D. **F.** Location of non-auditory ROIs. R: rostral; C: caudal.

**Figure 4.3:**
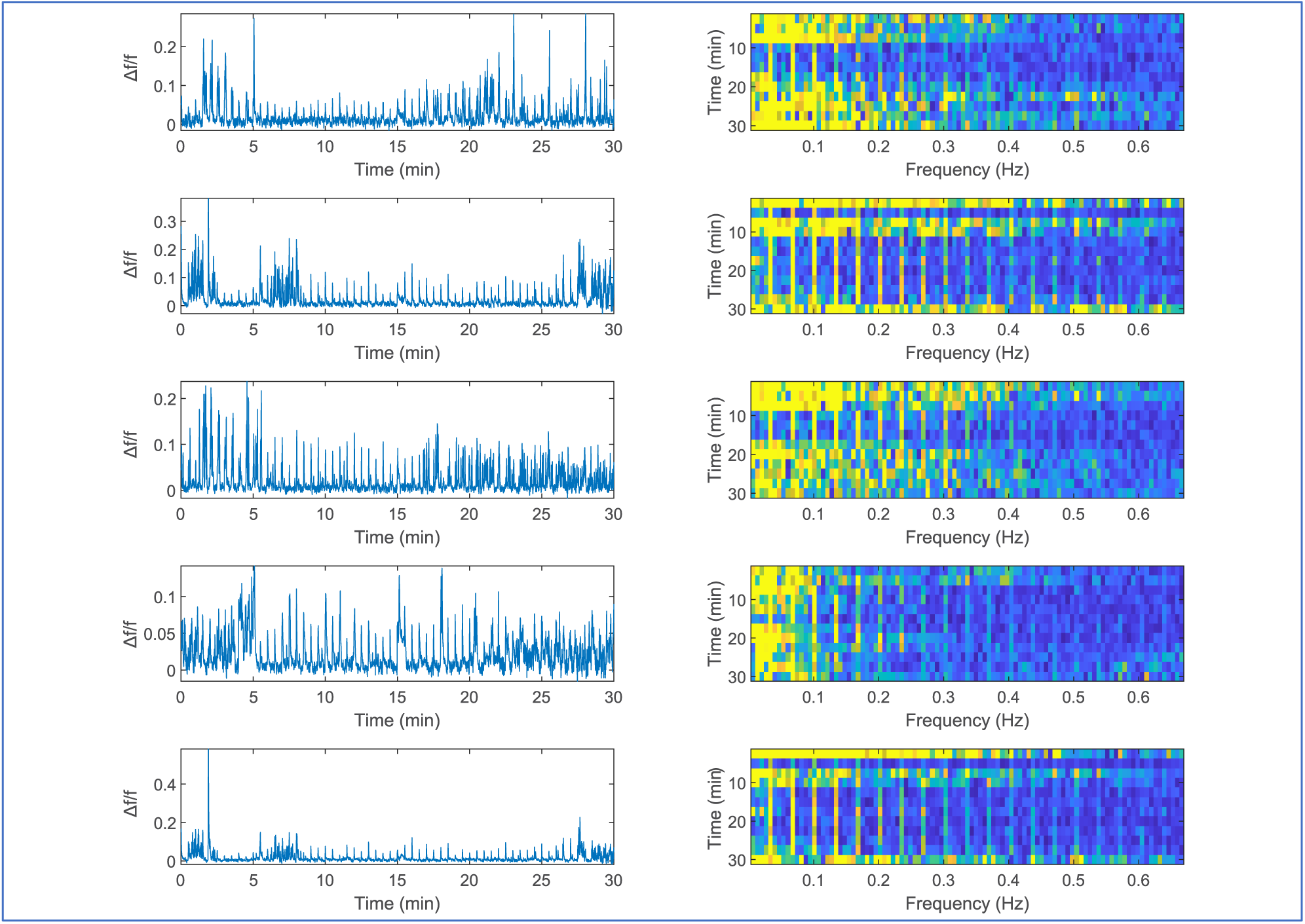
Examples of individual ROIs with decreased activity during DEX. Δf/f of 5 randomly chosen ROIs with decreased activity when exposed to DEX (left), drawn from the population of ROIs represented in Figure 4. Right: the same ROIs’ Fourier transforms for different time windows across the timeline.

**Figure 5.1:**
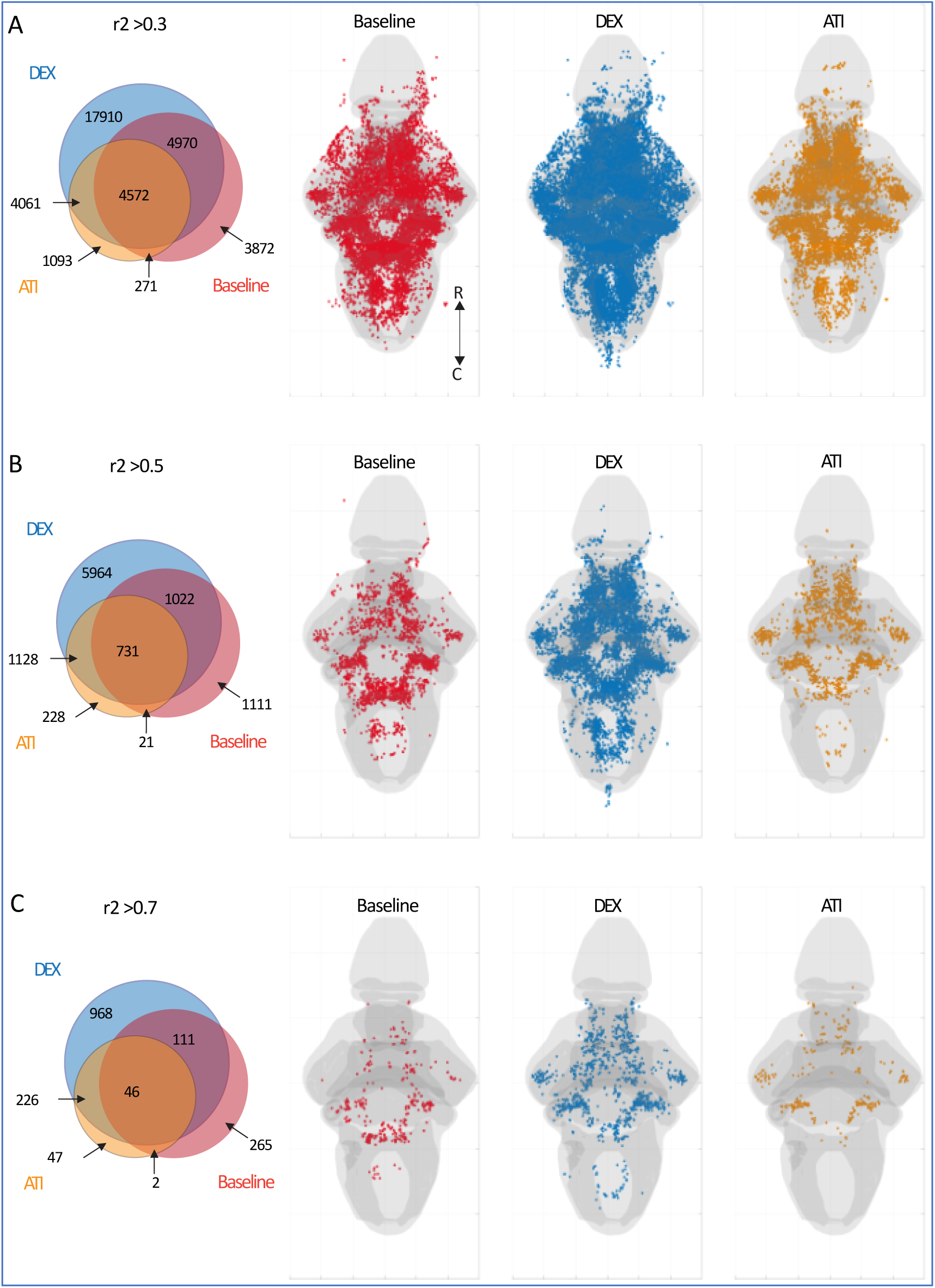
Detection of auditory neurons under DEX and ATI using linear regression. Venn diagram (left) showing the overlap of ROIs deemed to be auditory during baseline (red), DEX (blue), and ATI (yellow), and the corresponding spatial distributions of ROIs (right) from these populations Results are shown for different r^2^ thresholds: **A.** r^2^> 0.3. **B.** r^2^> 0.5. **C.** r^2^> 0.7.

